# Validation of a multiplex microsphere immunoassay for detection of *Trypanosoma cruzi* antibodies in dogs

**DOI:** 10.1101/2023.03.05.531211

**Authors:** Carlos A. Rodriguez, Rachel E. Busselman, Huifeng Shen, Ashley B. Saunders, Rick Tarleton, Sarah A. Hamer

## Abstract

The vector-borne protozoan parasite *Trypanosoma cruzi* causes Chagas disease in humans, dogs, and many other mammalian hosts. Canine Chagas disease is increasingly diagnosed in dogs of the southern US where triatomine insect vectors occur, and there are limited veterinary diagnostic options; currently, only the indirect fluorescent antibody (IFA) test is offered at a single accredited diagnostic laboratory. This study evaluated a multiplex microsphere immunoassay (MIA) for detecting antibodies against *T. cruzi* in dogs and compared with existing serological methods to establish cut-off values and relative sensitivity/specificity. Dog sera (n=135) which were previously characterized using the IFA and off-label use of two commercial rapid assays were tested on the multiplex MIA against 12 different antigens: nine *T. cruzi* antigens, a negative control recombinant protein (green fluorescent protein), a *Leishmania* antigen, and a canine parvovirus antigen (used as an antibody control given near-ubiquitous parvovirus vaccination). For each sample, the ratio of median fluorescence intensity (MFI) for each *T. cruzi* antigen to that of GFP was calculated. Samples with an antigen/GFP ratio greater than 4 standard deviations above the mean of 25 known negative sera were considered positive on that antigen. Samples testing positive on 2 or more antigens were considered positive for *T. cruzi* antibodies. Compared to the IFA, the multiplex MIA demonstrated a relative sensitivity of 100% and specificity of 96.97%. Given its precision, high-throughput format, potential for automation, and lack of subjective interpretation, the multiplex MIA should be considered a valid and improved assay for *T. cruzi* antibodies in dogs.

## Introduction

The vector-borne protozoan parasite *Trypanosoma cruzi* is endemic to the Americas and causes Chagas disease in a wide range of mammalian hosts, including humans and dogs. It is estimated that 288,000 people in the United States have Chagas disease, with origins of infection varying from travel-related to endemic spread^13^. Dogs are an important host of interest to veterinary public health, as they are an effective sentinel species and experience similar disease manifestations to humans^9, 19^. Previous studies in Texas have estimated the prevalence of Chagas disease in domestic dogs to be between 20.3% and 31.6%, with even higher estimates in kennel or shelter environments in Texas (up to 57.6%)^6, 7, 14^. The average yearly seropositivity in canine samples submitted to the Texas A&M Veterinary Medical Diagnostic Laboratory (TVMDL) is over 20%; additionally, demand for serologic testing at TVMDL has more than doubled in the last decade, from 1,129 test requests in 2014 to over 2,500 in 2022 (unpublished TVMDL data). Between 2014 and 2018, over 1,200 positive *T. cruzi* antibody tests were reported by TVMDL in samples submitted from Texas alone (unpublished TVMDL data).

Diagnosing Chagas disease can be challenging, as the parasite’s life cycle offers only brief windows of parasitemia during which *T. cruzi* organisms can be detected primarily by PCR of blood or by hematological methods. Evidence of the parasite in the heart and other affected tissues can be observed histologically, however this is primarily achieved post-mortem. As such, serologic methods are the primary means of antemortem testing for Chagas disease; since there is currently little evidence of patients self-clearing the parasite, seropositivity is typically considered a diagnosis of a current infection. In humans, the Centers for Disease Control and Prevention (CDC) considers a single test insufficient to definitively diagnose Chagas disease, and instead recommends using “two or more tests that use different techniques and detect antibodies to different antigens”^4^; this recommendation is also echoed by a recent publication on diagnosing Chagas disease in the United States^10^. For dogs, only the indirect fluorescent antibody (IFA) test is currently available as a validated option in accredited diagnostic laboratories; however, this approach requires specialized equipment, is interpreted using subjective evaluation of fluorescence patterns, and is prone to cross-reaction with other closely related pathogens (e.g., *Leishmania*) as it uses whole *T. cruzi* parasites as antigen. Commercially available rapid tests designed and approved for human use have been used in research settings to screen dogs for *T. cruzi* antibodies^17^, but this is not currently an option for routine diagnostics. Given these limitations, it is currently not practical to diagnose canine Chagas disease to the same level of rigor as the CDC’s recommendations for human diagnosis.

Human and veterinary diagnostic laboratories have begun developing and implementing multiplex microsphere immunoassays (MIA) for a variety of infectious diseases using a commercial platform known as xMAP Technology (Luminex Corp, Austin, TX)^20, 21^. These assays use microscopic polystyrene beads (available in magnetic or non-magnetic forms) as the solid phase, against which proteins of interest can be bound. For indirect assays, antibodies in the patient serum bind to the antigens on the beads, and a secondary antibody conjugated with phycoerythrin (PE) serves as a marker for any bound antibodies. Each bead is uniquely labeled with an internal dye, allowing the instrument reader to determine which target is represented while measuring the signal from the PE conjugate (indicating the presence/absence of antibodies in the patient serum against that target).

Previous studies^5, 12^ have described the use of a multiplex MIA to screen serum for *T. cruzi* antibodies in humans and dogs, determining a panel of appropriate antigens of interest and demonstrating effective diagnosis. Multiplex MIAs such as this one offer improvements over classical *T. cruzi* serology (i.e., IFA) in several ways: they eliminate the need for subjective interpretation by using an instrument reader; are performed using a 96-well plate, offering high throughput; increase precision via multiple readings per analyte, per sample; and offer high sensitivity and specificity by targeting multiple antigens. The aim of this study is to characterize this multiplex MIA and determine its validity in detecting antibodies against multiple *T. cruzi* antigens in dogs, establishing cut-off values for seropositivity and estimating sensitivity/specificity relative to existing methods. Additionally, we aim to demonstrate this assay’s utility in monitoring seropositive dogs over time.

## Materials and methods

Serum samples (n = 135) were previously collected from hunting dogs as part of a longitudinal study of Chagas disease in south Texas kennels, during which dogs were sampled at 0, 6, and 12 months^1^. A total of 61 individual dogs were represented in this sample set, comprised of 11 different breeds and ranging in age from 1 to 12 years old; 31 dogs were female, and 30 dogs were male. As part of the previous study, samples from these dogs were characterized using the IFA test and two commercially available rapid immunochromatographic tests (ICT); an endpoint titer was determined for the IFA by serially diluting samples until a final dilution with positive fluorescent signal was reached. The rapid ICTs (ChemBio StatPak and InBios Chagas Detect™ Plus) were used off-label to test dog serum, as they are both FDA licensed and validated for human use only. To determine and validate cutoff criteria for the MIA, we used a subset of samples that had congruent positive or congruent negative results across all 3 of the serological tests to which they were subjected. This included the samples which were categorized as clearly seropositive (n = 27) and clearly seronegative (n = 33) by the prior serological tests in the original study. Validating the assay with these samples of congruent serostatus was done because there remains uncertainty in the true infection status for animals that have discordant test results, and there is yet to be an accepted gold standard serologic test for *T. cruzi* exposure in human or veterinary medicine^8, 10, 15, 16^. Once cutoff criteria for the MIA were established with this sample set, we then used the full sample set which includes dogs sampled at multiple time points to monitor changes in serostatus over time.

A total of 12 antigens were evaluated in the multiplex assay (Table 1); 9 antigens from *T. cruzi*, were prepared and coupled to the xMAP beads as previously described^5^: FF10, G10, LE2, Kn107, FAB4, ATPase, Kn122, Kn80, and a whole-organism lysate. Green fluorescent protein (GFP) was used as a negative recombinant antigen. A canine parvovirus protein (VP2, MyBioSource, Inc., San Diego, CA)) was used as an antibody control; since all dogs in this study were assumed to have been previously vaccinated, samples failing to react to this protein would reflect inadequate antibody production or sample integrity issues. Lastly, a *Leishmania* protein (K39, MyBioSource, Inc., San Diego, CA) was included to examine cross-reactivity^18^. In addition to the study samples, 3 control sera were also be tested: negative and positive *Leishmania* controls, and a negative parvovirus control (VMRD, Inc., Pullman, WA).

**Table 1.** Proteins used as antigens in the canine *T. cruzi* multiplex MIA.

| Antigen name | Gene ID/description | Function in assay |
| --- | --- | --- |
| FF10 | TcCLB.506529.610 | *T. cruzi* antigen |
| G10 | TcCLB.504199.20 | *T. cruzi* antigen |
| LE2 | TcCLB.507447.19 | *T. cruzi* antigen |
| Kn107 | TcCLB.508355.250 | *T. cruzi* antigen |
| FAB4 | TcCLB.507029.30 | *T. cruzi* antigen |
| ATPase | TcCLB.509233.180 | *T. cruzi* antigen |
| Kn122 | TcCLB.506749.10 | *T. cruzi* antigen |
| Kn80 | TcCLB.506885.70 | *T. cruzi* antigen |
| Tc Lysate | *Trypanosoma cruzi* lysate from tissue culture | *T. cruzi* antigen |
| GFP | Green fluorescent protein | Negative control recombinant protein |
| VP2 | Canine parvovirus capsid protein VP2 | Positive antibody control protein |
| K39 | Kinesin-like protein, partial | *Leishmania* antigen |

Samples were tested as previously described for dog serum^12^. A mixture containing all 12 antigen-coupled beads in assay buffer (PBS with casein) was prepared and added to 96-well flat bottom plates. The plates were affixed to magnetic holders and the beads allowed to settle at the bottom of the wells. The excess assay buffer was removed from the plates, leaving the beads behind, and 100 μL of the serum samples (diluted 1:500) added in duplicate wells. One set of wells had only plain assay buffer added as a blank/background sample. The plates were sealed with foil and incubated at room temperature on a shaker for 1 hour. At the end of the first incubation, the plates were washed three times; for each wash, the magnetic holder was used to keep the beads in place, and 200 μL of assay buffer was added and then removed from the wells. Anti-canine IgG conjugated with phycoerythrin (Rockland, Inc., Limerick, PA) was used as a secondary antibody at a working dilution of 1:100; 150 μL of the PE conjugate was added to each well, the plates again sealed with foil, and incubated at room temperature on a shaker for 1 hour and 30 minutes. After the second incubation, the plates were again washed three times. The beads were then resuspended with 150 μL of assay buffer and read using a MAGPIX instrument equipped with xPONENT software (Luminex Corp, Austin, TX), and the median fluorescence intensity (MFI) for all 12 antigens in each well was measured. The average net MFI (average MFI of both sample wells minus the MFI of the blank/background wells) for each antigen on each sample was used for subsequent analysis.

For all samples, the average net MFI of each antigen was divided by that of GFP (hereon referred to as the antigen-GFP ratio). This allowed analysis to be performed on a normalized value as well as corrected for any samples which had high reactivity against GFP (i.e., background or non-specific reactions)^22^. A subset of samples (n = 25) which were previously negative for *T. cruzi* antibodies by all 3 existing serologic methods were designated as a “negative pool”, and the mean antigen-GFP ratio for each antigen was calculated for these samples. Study samples with antigen-GFP ratios above 4 standard deviations from the mean of the negative pool were considered reactive to that antigen. Statistical analyses were performed using Stata software (StataCorp LLC, College Station, TX), and graphical representations of results via heatmaps were generated using Prism 9 software (GraphPad Software, La Jolla, CA) to compare results between groups and visualize seroconversion over time. A receiver operating characteristic (ROC) using the IFA as the reference test was used to determine the optimal number of reactive antigens required to designate a sample as positive for antibodies against *T. cruzi*, along with assay sensitivity/specificity relative to the IFA. Correlation analysis was performed to investigate the relationship between IFA endpoint titer and the number of positive MIA antigens.

## Results

The area under the curve (AUC) for the ROC was 0.9989 (Figure 1), indicating the multiplex assay is highly comparable to the reference test (IFA). The ROC analysis indicated an optimal cutoff of ≥2 reactive antigens to classify a sample as seropositive (Table 2). Using this criterion, the multiplex assay had relative sensitivity of 100% and specificity of 96.97%, with only a single result disagreeing with the reference test. The group of samples which were positive on IFA had notably higher antigen-GFP ratios than the IFA negative group (Figure 2). The number of reactive antigens for seropositive samples was highly correlated with the previous IFA titer (r = 0.8675, p<0.0001). All samples were reactive on the parvovirus protein, indicating no apparent antibody failures or sample quality issues. The *Leishmania* antigen was functional, with the positive control showing high reactivity and the negative control showing none (Figure 2). Notably, the *Leishmania* positive control was also reactive on 2 *T. cruzi* antigens (whole-organism lysate and Kn107). A single *T. cruz-*negative sample was considered reactive for the *Leishmania* antigen using the diagnostic criteria established for *T. cruzi* (4SD above the *T. cruzi*-negative pool).

**Figure 1.**
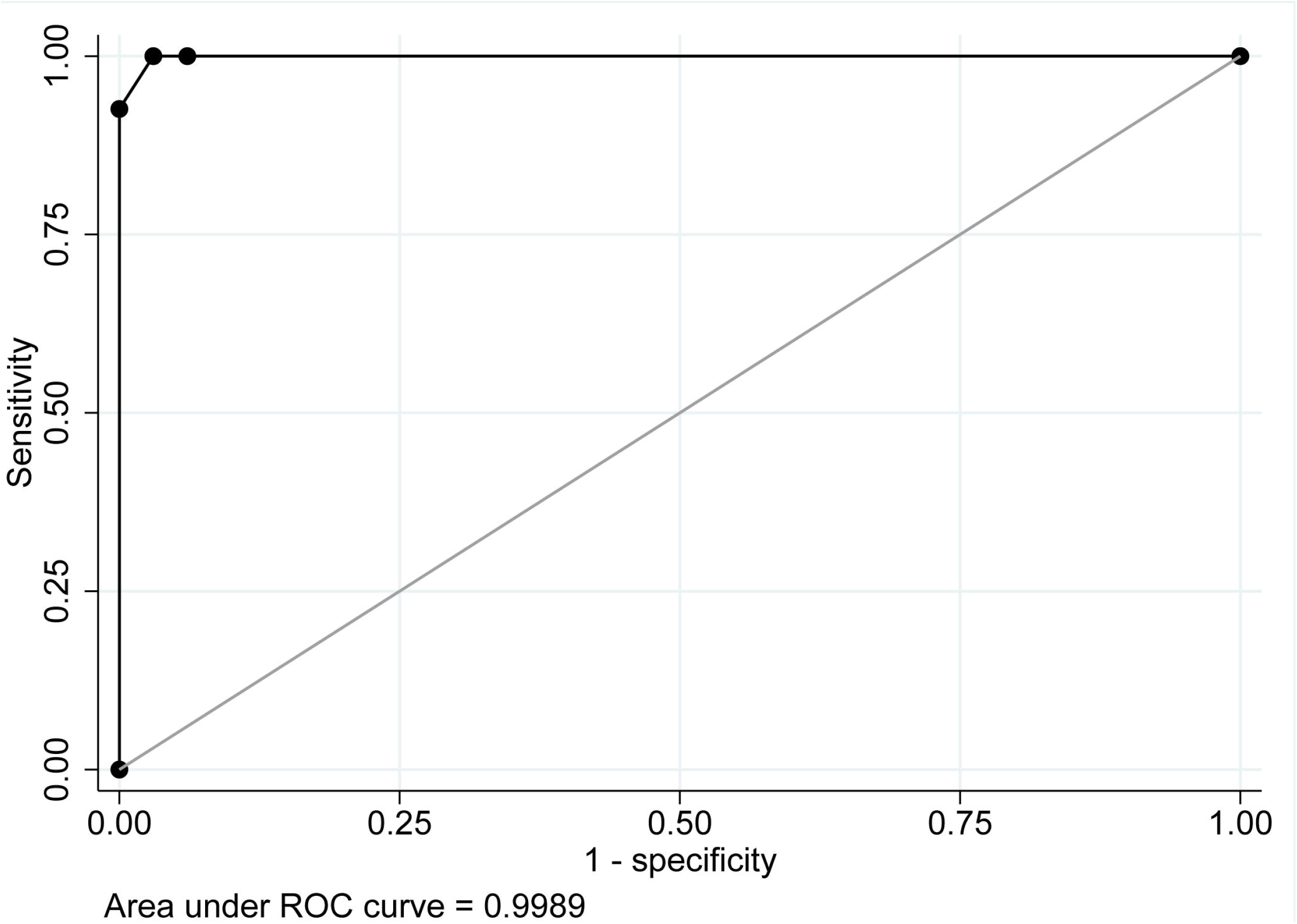
Receiver operating characteristic (ROC) curve for the canine *T. cruzi* antibody MIA using the IFA as a reference standard.

**Table 2.** Sensitivity, specificity, and percentage of samples correctly classified at different cut-off values for the canine *T. cruzi* multiplex MIA relative to the IFA.

| Cut-off<br>(# positive antigens) | Sensitivity | Specificity | % Correctly classified |
| --- | --- | --- | --- |
| ≥ 1 | 100.00% | 93.94% | 96.67% |
| ≥ 2 | 100.00% | 96.97% | 98.33% |
| ≥ 3 | 92.59% | 100.00% | 96.37% |
| ≥ 4 | 48.15% | 100.00% | 76.67% |
| ≥ 5 | 11.11% | 100.00% | 60.00% |
| ≥ 6 | 3.70% | 100.00% | 56.67% |

**Figure 2.**
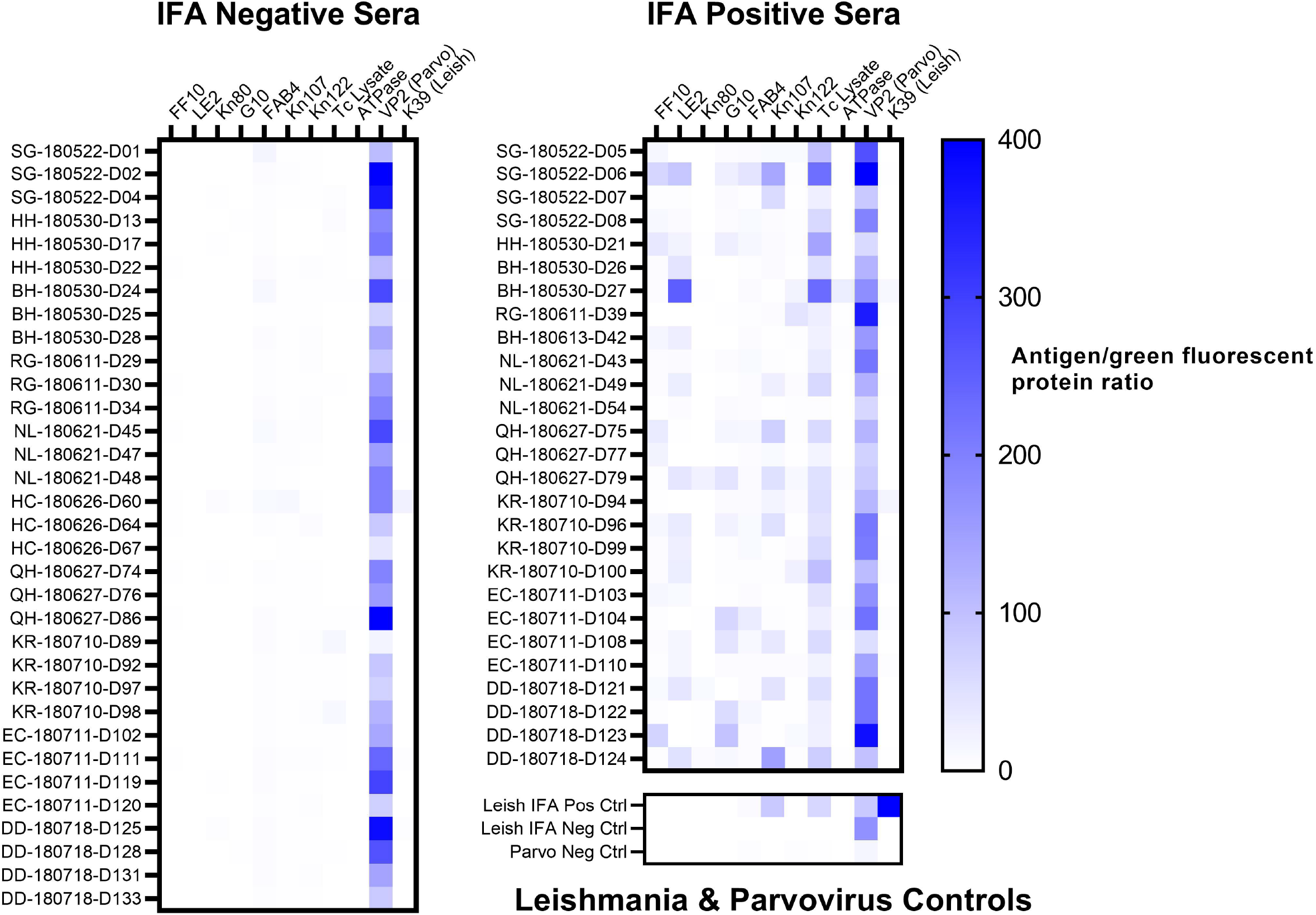
Heatmap showing reactivity of 60 canine serum previously tested for *T. cruzi* antibodies by IFA (grouped by previous IFA result), and reactivity of *Leishmania* & canine parvovirus controls.

Of the 37 dogs for which all three longitudinal samples were tested (Figure 3), 8 were considered in the original study to have seroconverted between sampling points based on a consensus result of at least 2 positive serologic tests^1^. Using the MIA, 3 (37.5%) of these dogs showed apparent seroconversion using the optimized diagnostic cut-off values used in this study (2 or more positive antigens); 2 more dogs were below the cut-off criteria but were seropositive against a single antigen. Once seropositive, the number of reactive antigens remained relatively stable over time, fluctuating on no more than 2 antigens for most dogs (and with no dogs reverting from seropositive to seronegative over time). All samples in the longitudinal group which were previously positive by IFA were positive on the multiplex assay, and the patterns and magnitude of reactive antigens were unique between individual dogs.

**Figure 3.**
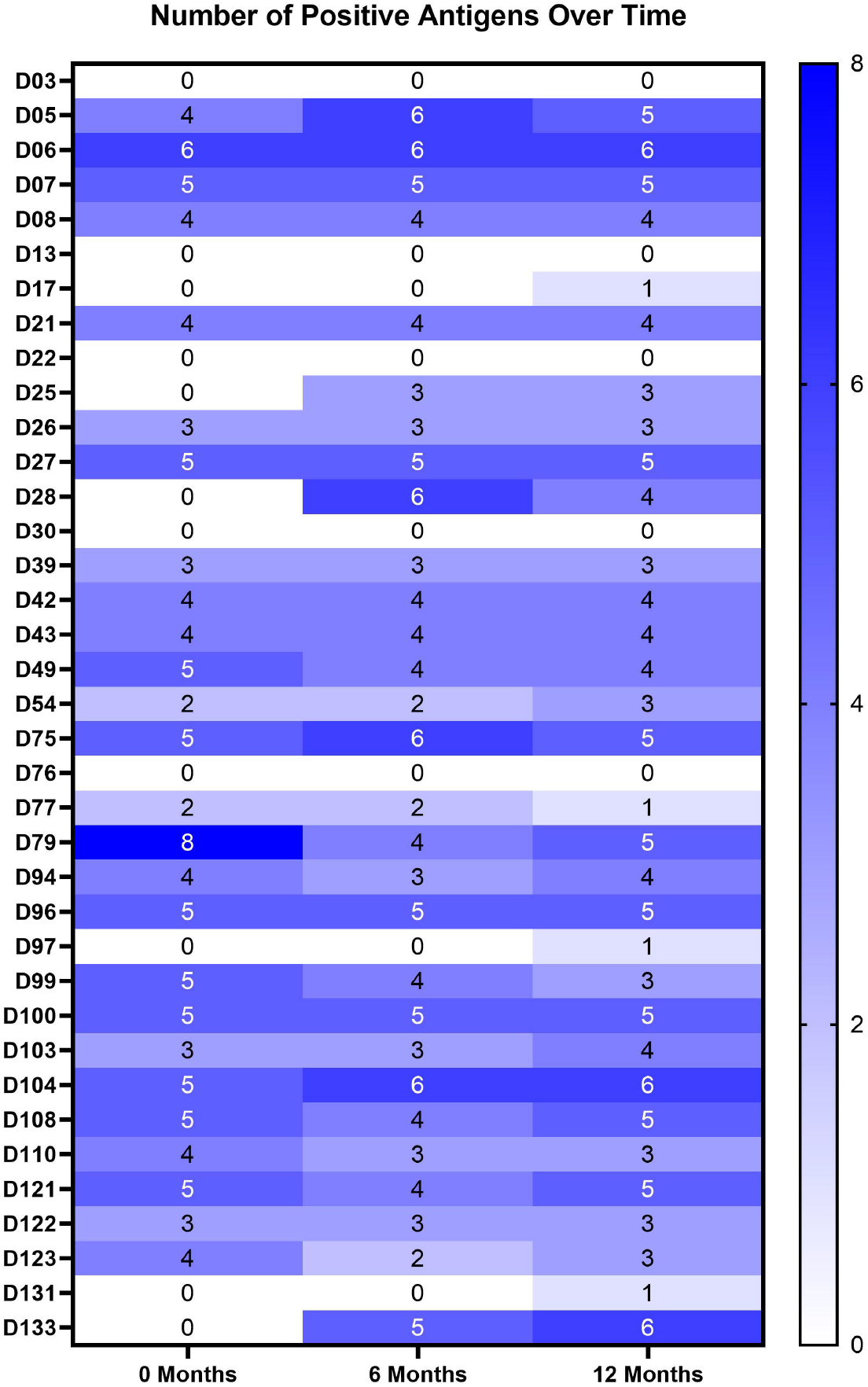
Heatmap showing the number of positive *T. cruzi* antigens on the MIA at 0-, 6-, and 12-month intervals for 37 dogs presumed to be positive based on previous testing.

## Discussion

Our study successfully demonstrated the utility of a multiplex microsphere immunoassay in detecting antibodies against *T. cruzi* in dogs when compared to existing test methods. Although other serological methods such as IFA and rapid immunoassays have been used extensively in diagnostic and research settings and are considered validated for those purposes based on human literature and correlation with clinical signs, few previous studies have been conducted which quantify their performance in terms of sensitivity and specificity relative to a gold standard. The multiplex MIA has been used in previous studies to test dogs, humans, non-human primates, and mice^3, 5, 12^; however, this is the first study to estimate its sensitivity/specificity for diagnostic purposes. While this study determined an optimal single cut-off point of 2 positive antigens to balance the inverse relationship between sensitivity and specificity, when implementing this assay in a true diagnostic setting, it may be most useful to utilize a three-tiered approach bracketing cut-off values to maximize both values (i.e., classifying samples as negative below the cut-off which gives 100% sensitivity, positive at the cut-off which gives 100% specificity, and “suspect” in between).

Evaluation of the multiplex MIA’s performance was dependent on several assumptions. First, although there is no perfect “gold standard” for *T. cruzi* testing, the IFA was used as the reference test in the ROC analysis and determination of sensitivity/specificity for the MIA. Second, antibody positive dogs were assumed to be infected; this is due to the lack of evidence of self-clearing once an animal has been exposed to the *T. cruzi* parasite. The sample size in this study was also relatively small due to the decision to use well-characterized samples. Additionally, the samples selected for validation were those which had the clearest seropositive/negative reactions based on previous testing, and our study did not explore the MIA’s performance in classifying discordant or “borderline” results in animals which are truly infected/not infected; this may positively bias the estimates of sensitivity/specificity in our study^11^. Future studies could improve on our estimates of Se/Sp using larger sample sets and statistical analyses which account for the lack of a gold standard (e.g., latent class analysis). Regardless of these limitations, the multiplex assay demonstrated high agreement with the IFA, and could perhaps serve as a new and improved “gold standard” going forward given its improvements over the more classical method.

Existing *T. cruzi* antibody assays are widely known to cross-react with *Leishmania* antibodies, confounding diagnosis for animals who could plausibly have been exposed to both pathogens. In addition to a whole-organism lysate, the inclusion of individual *T. cruzi* proteins in this assay (including at least one—FF10—which is completely unique to this organism) is intended to increase specificity. Given that the *Leishmania* positive serum was positive on the *T. cruzi* assay, but none of the *T. cruzi* positive samples were significantly reactive on the *Leishmania* antigen, there is likely not enough evidence in this study to conclude whether cross-reactivity will carry over to this testing platform. Additional studies with known *Leishmania* positive and negative samples, from areas in which *T. cruzi* exposure is unlikely, could ultimately provide more definitive answers to this question. Previous studies^2, 3^ have used different sets of *T. cruzi* proteins in the multiplex assay, and one of the advantages of this platform is the ability to expand/modify antigens to best optimize the test. If implementing this assay in settings where more definitive discrimination from *Leishmania* positivity is crucial (e.g., where infection with either pathogen is a possibility), additional proteins which are completely unique to *T. cruzi* may be added to increase specificity.

When compared to the longitudinal results of the Busselman et al. study, the MIA showed perfect agreement with the IFA results, but did not detect seroconversion against any antigens in 3 of the 8 dogs which had apparently seroconverted based on positive results on 2 rapid tests (Figure 3). As these dogs were consistently negative by both IFA and MIA, it is possible that the rapid tests are more sensitive (and thus less specific) than either of these methods. The 2 dogs which appeared to seroconvert against only 1 MIA antigen did so only during their final sampling point; of note, these dogs were also PCR positive at this sampling point. Although these dogs were originally considered seronegative according to the diagnostic criteria of 2 or more positive antigens, this may be an indication that positivity on even a single antigen can indicate early infection and should be considered as a suspicious result in within the clinical and epidemiological context. As such, the multiplex assay may be useful in monitoring at-risk populations of dogs (e.g., in kennel settings) over time, where a change in seropositivity between sampling points—even on a single antigen—could trigger immediate follow-up testing by PCR, allowing diagnosis and possible intervention for acute cases sooner than conventional serology alone.

## Acknowledgements

We thank Lisa Auckland for assistance with sample preparation. We thank Dr. Pam Ferro and TVMDL for support throughout this project. Negative canine parvovirus serum was kindly provided by Ed Felt and VMRD, Inc. We also thank Heather Darby and Luminex Corporation for on-site training and support in setting up the MAGPIX instrument.

## Declaration of conflicting interests

The authors declared no potential conflicts of interest with respect to the research, authorship, and/or publication of this article.

## Funding

American Kennel Club Canine Health Foundation Grant No 02448 provided funding for collection of the dog samples used in the study.

## Notes

### Competing Interest Statement

The authors have declared no competing interest.

